# ShinyGO: a graphical enrichment tool for ani-mals and plants

**DOI:** 10.1101/315150

**Authors:** Steven Xijin Ge, Dongmin Jung

## Abstract

**Motivation:** Gene lists are routinely produced from various genome-wide studies. Enrichment analysis can link these gene lists with underlying molecular pathways by using functional categories such as gene ontology (GO).

**Results:** To complement existing tools, we developed ShinyGO with several features: (1) large annotation database from GO and many other sources for over 200 plant and animal species, (2) graphical visualization of enrichment results and gene characteristics, and (3) application program interface (API) access to KEGG and STRING for the retrieval of pathway diagrams and protein-protein interaction networks. ShinyGO is an intuitive, graphical web application that can help researchers gain actionable insights from gene lists.

**Availability:** http://ge-lab.org/go/

**Supplementary information:** Supplementary data are available at *Bioinformatics* online.

## 1 Introduction

For a set of genes identified in genome-wide studies, enrichment analysis can be done to see if it is enriched with genes of a certain pathway or functional category. One of the most widely used functional categories is the gene ontology (GO) (Ashburner, et al., 2000). Due to its importance for the interpretation of gene lists, dozens of tools have been developed (Khatri, et al., 2012).

A small subset of these tools is listed in Supplementary Table S1 in supplementary file 1. Some tools are designed for biomedical research and thus focus primarily on human and mouse. For example, Enrichr (Kuleshov, et al., 2016) includes a comprehensive database of 234,849 annotated gene-sets in 128 libraries, ranging from GO, co-expression, tis-sue-specific genes, transcriptional factor (TF) or microRNA (miRNA) target genes, to various pathway databases. Similarly, tools like PlantGSEA are focused on 15 plants species(Yi, et al., 2013).

g:Profiler is based on gene annotation in Ensembl (Aken, et al., 2017) for over 200 plant and animal species. STRING (Szklarczyk, et al., 2015) is large database of protein-protein interactions (PPI). It also provides functionality for enrichment analysis of GO and protein domains in 115 archaeal, 1678 bacterial, and 238 eukaryotic species (version 10). On the other extreme is DAVID (Huang da, et al., 2009) which covers tens of thousands of organisms. This is possible because of the it derives information from many sources including NCBI, UniProt, KEGG, GO, Biocarta, REACTOME, etc. These tools have help biologists gain insights from gene lists.

In the current study, we develop a new tool based on the relatively large annotation database at Ensembl. Compared with existing tools, ShinyGO has improved graphical visualization of enrichment results and the ability to display pathway diagrams and protein interaction networks.

## 2 Methods

ShinyGO is a Shiny application developed based on several R/Bioconductor packages, and a large annotation and pathway database compiled from many sources. See Supplementary file 1 for more details. Source code is available at https://github.com/iDEP-SDSU/idep/tree/master/shinyapps/go.

## 3 Results

We developed ShinyGO for in-depth analysis of gene lists, with graphical visualization of enrichment, pathway, gene characteristics, and protein interactions (Figure 1). It is based on annotation databases for 208 organisms, including 97 at Ensembl (vertebrates, release 91) (Aken, et al., 2017), 45 from Ensembl Plants (release 37) (Bolser, et al., 2017), and 66 from Ensembl Metazoa (release 37). See supplementary file 2 for a list. We batch downloaded not only GO functional categorizations, but also gene ID mappings and other quantitative gene characteristics. Query genes are mapped to all gene IDs in the database, for both ID conversion and suggestion of possible organisms.

**Figure 1.**
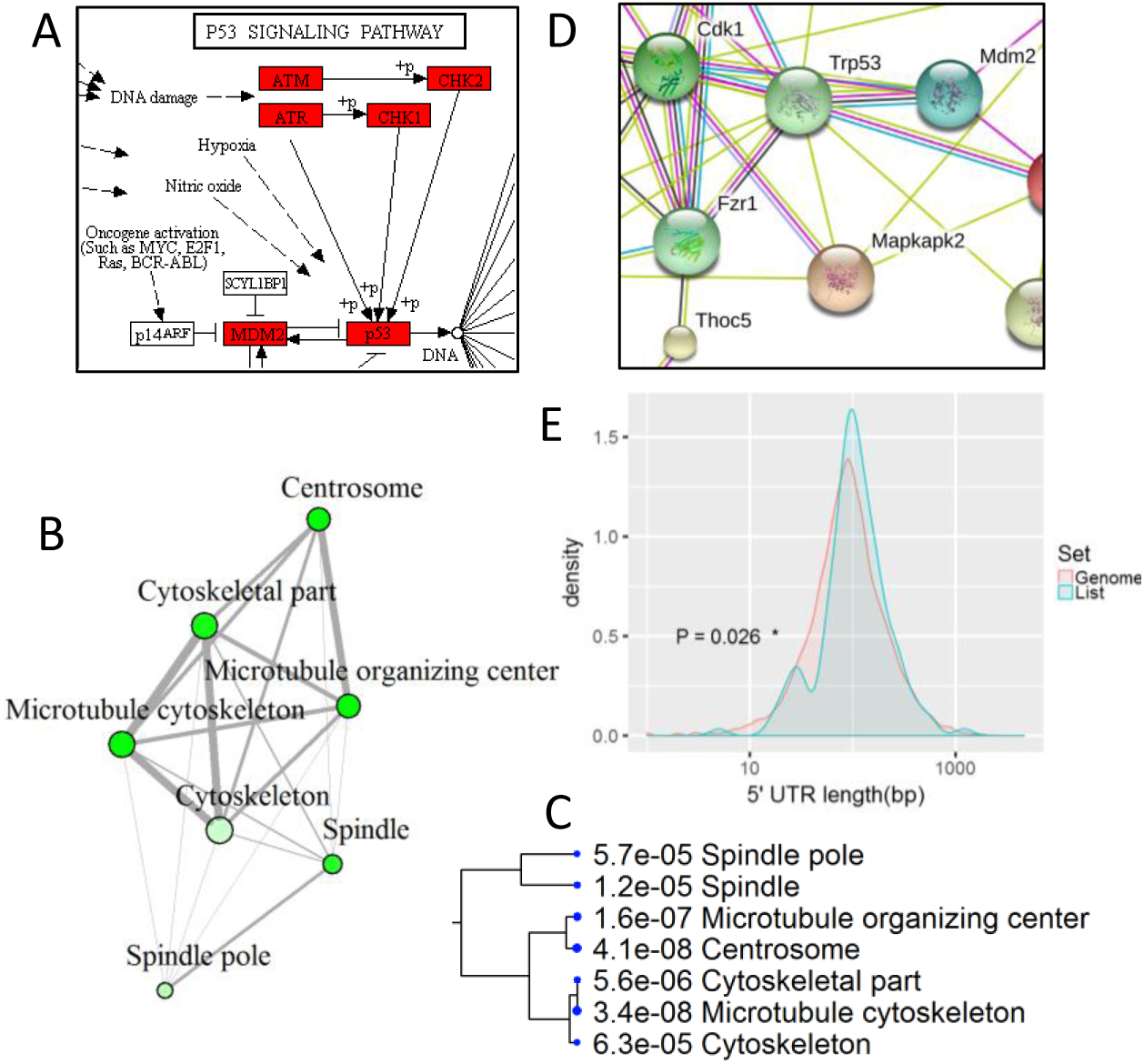
Example outputs of ShinyGO. A). A partial KEGG pathway diagrams with genes highlighted. Enriched GO molecular component terms visualized as a network (B) and hierarchical clustering tree(C). D). PPI network. E. Distribution of the lengths of 5’ UTRs in query genes vs. other coding genes in the genome.

In addition to GO, pathways were downloaded directly from KEGG (Kanehisa, et al., 2017). For human, various pathway data are also obtained from MSigDB (Liberzon, et al., 2015), GeneSetDB (Araki, et al., 2012), Reactome (Fabregat, et al., 2016), as well as many sources of verified or predicted miRNA and TF target genes. Totally, we compiled 72,394 gene-sets for human (Supp. Table 2). Similar databases, such as GSKB (Lai, 2016) for mouse and araPath (Lai, et al., 2012) for Arabidopsis, are included in ShinyGO.

ShinyGO can retrieve pathways diagram from KEGG web server via API access using the pathview Bioconductor package (Fig. 1A). To visualize overlapping relationships among enriched gene-sets, we developed a network view (Fig. 1B) and a tree view (Fig. 1C) of the enriched gene-sets. In a GO cellular component (CC) enrichment analysis, Fig. 1B and 1C shows that three terms related to cytoskeleton overlaps in many genes.

ShinyGO also plots the chromosomal locations of all the genes in the user’s list. It also detects whether the genes are randomly distributed on the chromosomes using a Chi-squared test, compared with other genes in the genome. Similar tests are also conducted to see if query genes differ from the rest in terms of the number of exons and transcript isoforms, and the types of genes (coding, non-coding, pseudogenes and so on). We plot the distribution of GC content, and the lengths of coding sequences, transcripts, and UTRs (untranslated regions). T-tests are carried out to identify any significant differences. As shown in Figure 1E, the query genes seem to have longer 5’ UTRs than other genes in the genome.

Enrichment analysis can also be conducted through API access to STRING (Szklarczyk, et al., 2015), thus expanding the number of covered organisms. PPI networks are retrieved directly from STRING. ShinyGO also produces a custom link to an interactive, annotated network on the STRING web site (Fig. 1D) with protein structures and Pubmed.

A use case can be found in supplementary file 1 with many example outputs. Through the analysis of 147 human genes upregulated by radiation, we were able to identify some expected pathways such as p53-mediated DNA damage response, as well as the underlying TFs (p53 and Rela/NF-κB) and even miRNAs ( miR-145 and miR-21).

## 4 Discussions

ShinyGO is an intuitive, graphical tool for enrichment analysis. Even though its species coverage is not a broad as DAVID, ShinyGO has more comprehensive gene-sets regarding TF and miRNA target genes for human, mouse and Arabidopsis. We will continue to compile such information for other organisms and update the annotation database on a yearly basis. To improve reproducibility, we plan to make older versions of the database available to users.

## Acknowledgements

The authors thank En Woo Sun, Brian Moore, and Kevin Brandt for technical support.

